# Leveraging vision to understand curiosity

**DOI:** 10.1101/2022.09.23.509220

**Authors:** Michael Cohanpour, Mariam Aly, Jacqueline Gottlieb

## Abstract

Humans are immensely curious and motivated to reduce uncertainty. Inspired by studies of sensory uncertainty, we hypothesized that visual areas provide multivariate representations of uncertainty, which are read out by higher-order areas that encode confidence and, ultimately, translated into curiosity. During fMRI, participants rated their confidence in identifying distorted images of animals and objects and their curiosity to see the clear image. We found that curiosity peaked at low confidence. To link sensory certainty and curiosity, we estimated “OTC Certainty” as the product of absolute and relative evidence for animals vs. object categories in occipitotemporal cortex (OTC) while participants viewed the distorted image. OTC Certainty negatively correlated with curiosity, while univariate activity in two frontal regions – vmPFC and ACC – positively correlated with confidence. The relationship between OTC certainty and curiosity was mediated by the vmPFC but not ACC. The results reveal neural mechanisms that generate curiosity from representations of uncertainty.

## Introduction

Humans are immensely curious – we are motivated to seek information even if this comes with no monetary rewards (Gottlieb & Oudeyer, 2018; Kidd & Hayden, 2015; van Lieshout et al. 2020). Curiosity compels us to explore, inquire, learn, and discover. What neural mechanisms underlie curiosity?

Recent studies of curiosity have used tasks in which participants rate their curiosity about trivia questions (Kang et al. 2009; Gruber et al. 2014; Baranes et al. 2015). These studies have found that curiosity ratings are encoded in brain regions implicated in motivation and reward, suggesting that systems that encode the value of primary reinforcers also encode the higher-order value of information (Kang et al. 2009; Gruber et al. 2014). Moreover, curiosity about trivia questions spurs cognitive actions geared toward processing the obtained information, including anticipatory attentional shifts toward the location of the answer (Baranes et al. 2015) and enhanced memory for the answer mediated by functional interactions between the hippocampus and reward circuitry (Gruber et al. 2014; Murphy et al 2021).

Despite these important advances, challenging open questions remain about the mechanisms that generate curiosity: how does the neural representation of an event lead to feelings of curiosity about that event? Theoretical models propose that this link involves a participant’s confidence about an event (Loewenstein 1994; Golman & Loewenstein 2018; Berlyne 1954), as low or intermediate confidence evokes high curiosity (Kang et al. 2009; Baranes et al. 2015). However, the neural underpinnings of this hypothesis have been difficult to evaluate, because little is known about the neural representations of confidence or certainty about trivia questions. Here, we develop a new task of perceptual curiosity -- the desire for information about ambiguous or distorted sensory stimuli (Berlyne, 1954; Nicki et al. 1970; Jepma et al. 2012) – that allows us to examine this question and link curiosity with the neuroscience literature on visual representations, confidence, and uncertainty.

The neural basis of confidence has been studied in perceptual tasks in which participants report decisions about visual stimuli as well as their confidence about properties of the stimuli. These studies have shown that confidence ratings depend on both sensory features and behavioral factors like response heuristics, response bias, or context (Maniscalco et al. 2016; Peters et al. 2017; Zylverberg et al. 2012; Merkel et al. 2008). Moreover, converging empirical evidence from studies using functional magnetic resonance imaging (fMRI), lesions, experimental manipulations, and inter-individual variability shows that confidence ratings depend critically on the frontal lobe (Lebreton et al 2015; Gherman et al. 2018; Del Cul et al. 2009; Rounis et al. 2010; Fleming et al. 2010; Fleming et al. 2014), consistent with the broader role of this area in metacognitive evaluation (Shimamura 2000; Fleming & Dolan 2010).

In addition, distinct neural signals of certainty are thought to be embedded in sensory representations. Evidence from monkey neurophysiology (Ma et al. 2006; Walker et al. 2020), human brain imaging (van Bergen et al. 2020; Geurts et al. 2022), and computational modeling (Ma et al 2008, Meyniel et al. 2015) suggests that feature-tuned neurons in early visual cortex simultaneously represent the most likely visual feature and the certainty about the presence and identity of those features. Importantly, while frontal cortical responses encode confidence in a *univariate* fashion (i.e., through overall changes in average fMRI BOLD response; Lebreton et al 2015; Gherman et al. 2018), certainty in sensory areas is conveyed by patterns of activity across neurons or voxels selective to visual features, and has been referred to as *multivariate certainty* (Russell et al. 2019).

Although much remains to be learned about how univariate confidence and multivariate certainty are related, computational models propose that multivariate representations of certainty are read out by associative areas that univariately encode confidence (Meynial et al. 2015; Pouget et al. 2016). This view is consistent with the representational untangling hypothesis, whereby multivariate neural representations are transformed into simpler, lower-dimensional representations that can be more readily used for controlling behavior (DiCarlo & Cox 2007; Russo et al. 2018). Empirical support for this theoretical view comes from a recent report showing that the confidence of human observers in a task of orientation discrimination is related to the multivariate certainty decoded from primary visual cortex using fMRI (Geurts et al 2022).

A key open question, however, is whether and how this distributed circuitry relates to curiosity. The handful of studies that probed perceptual curiosity did not simultaneously require ratings of confidence and curiosity, measure brain activity, or quantify multivariate certainty (Nicki et al. 1970; Jepma et al. 2013). Moreover, while these studies have shown that perceptual curiosity is triggered by complex natural stimuli (Nicki et al. 1970; Jepma et al. 2013), we lack methods that can decode multivariate certainty regarding such stimuli. To date, the studies investigating multivariate certainty have focused exclusively on elementary visual features (e.g. orientation; Walker et al. 2020; Geurts et al. 2021). For example, the algorithm that Geurts et al. developed for decoding multivariate certainty from human V1 depends critically on the assumption that orientation is encoded by neurons with well-defined cosine tuning curves (vanBergen et al. 2021; Geurts et al 2022). In contrast, natural stimuli that evoke curiosity are represented in higher-level visual areas in which single-neuron tuning is poorly defined. One such area is the occipitotemporal cortex (OTC) which is crucial for object recognition (Grill-Spector & Malach, 2004; Kar et al. 2019) and encodes biologically relevant stimulus categories (e.g., animals vs man-made objects) in its multi-voxel activity patterns (Long et al. 2018; Konkle & Oliva 2012; Konkle & Caramazza 2013). However, no study has quantified multivariate certainty of these higher-level representations.

To examine these questions, here we used a new behavioral task in which we elicited perceptual curiosity using synthetic images of animals and man-made objects that were distorted according to a well-defined algorithm (“texforms”) and designed to activate OTC multi-voxel category representations (Long et al. 2018). We developed a trial-by-trial metric of the multivariate certainty in OTC multi-voxel patterns (“OTC Certainty”) that avoided restrictive assumption of canonical tuning curves (vanBergen et al. 2021; Geurts et al. 2022) and was inspired instead by the modern machine learning literature (Huellermeir & Weigeman 2020). Finally, we examined how OTC certainty was related to behavioral ratings of curiosity and univariate activity in two frontal areas implicated in perceptual confidence: the ventromedial prefrontal cortex (vmPFC), which has direct anatomical connections with the OTC (Catanati 2005; Furl 2015) and encodes visual confidence in multiple tasks (Gherman et al. 2018; Hebscher et al. 2016; Lebreton et al. 2015), and the anterior cingulate cortex (ACC), which is critical for cognitive control (Shenhav et al. 2013) and is implicated in visual confidence and curiosity (Bang et al. 2018; Geurts et al 2022; Jepma et al 2013).

Similar to curiosity about trivia questions, perceptual curiosity has a negative, quadratic relationship with confidence (Gruber et al. 2014; Baranes et al. 2015). Most importantly, OTC Certainty negatively correlates with perceptual curiosity and this correlation is mediated by univariate representations of confidence in the vmPFC but not the ACC. The results link perceptual curiosity with neural representations of confidence and certainty and shed light on the mechanisms by which the neural representation of an event generates curiosity about that event.

## Results

### The perceptual curiosity task

Thirty-two participants (17 female) completed a task (*Methods, Design & Procedure, Perceptual Curiosity Task*) that measured perceptual curiosity while undergoing fMRI scanning. To evoke perceptual curiosity, we presented participants with texforms (**Fig. 1A**), distorted images that preserve some texture and form information but are difficult to recognize at the exemplar level (Long et al. 2018). We used texforms of animals and man-made objects so that we could reliably activate OTC multivoxel patterns. On each trial, participants viewed a texform of an animal or man-made object and were instructed to imagine their best guess for what the original (undistorted) image was, and to rate their confidence in their best guess and their curiosity to see the original image (**Fig. 1B**). After their ratings, participants were shown the original image. This procedure is similar to studies of epistemic curiosity (Kang et al. 2009; Gruber et al. 2014; Baranes et al. 2015) except that participants reported their curiosity and confidence about a visual image rather than a trivia question; thus, the texforms can be considered the “questions” and the undistorted image the “answers” in our task. Participants reported all the ratings by rotating an MRI-safe trackball to position a cursor on a 0-100 scale, with the initial cursor position randomized to control for motor confounds. Each participant completed 84 trials divided evenly into four runs. To ensure that the ratings were unbiased by instrumental incentives, participants received a fixed compensation for completing the task ($40) but no payoffs based on the ratings they gave.

**Figure 1:**
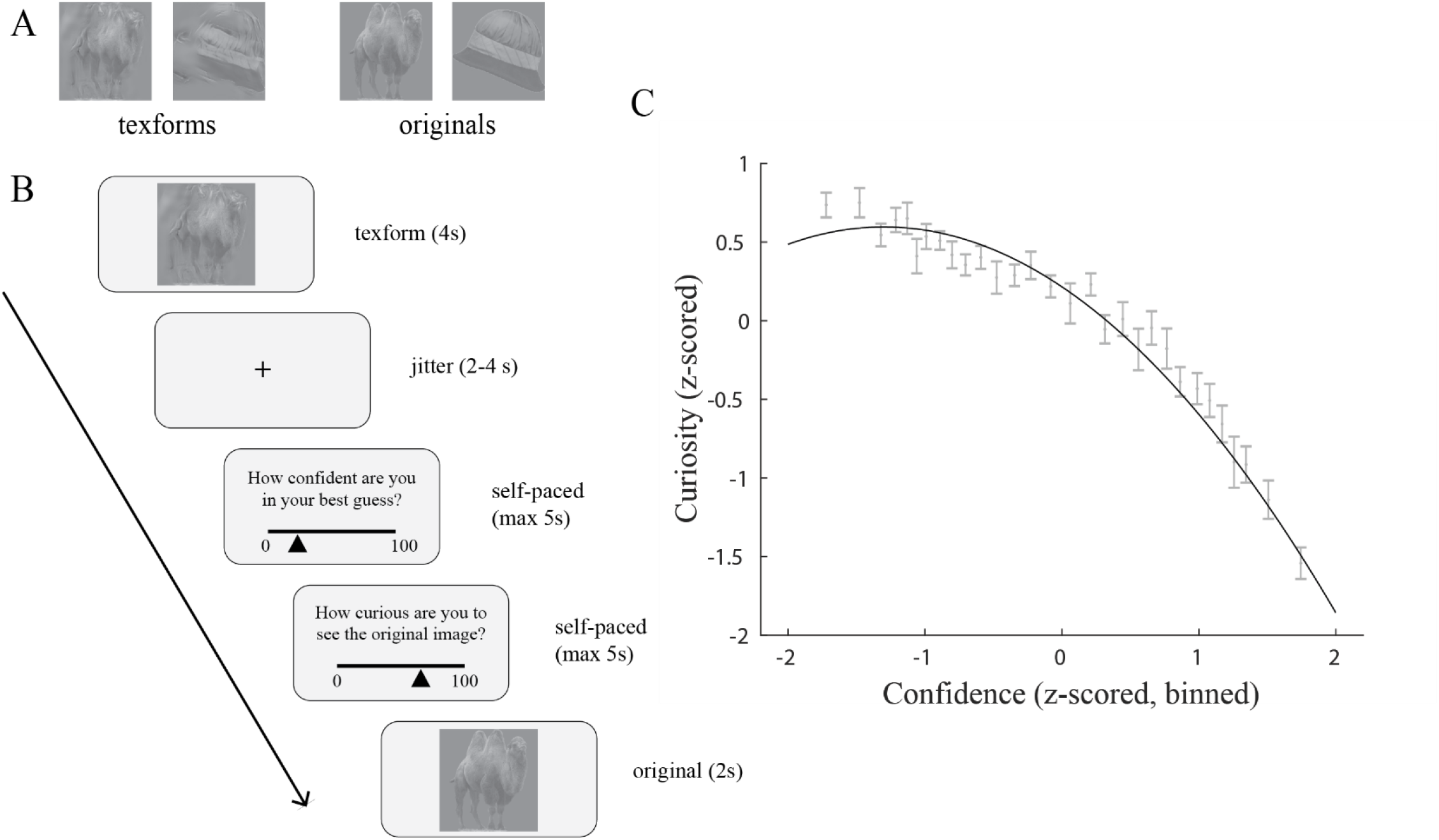
(A) **Stimuli**. Texforms of camel and hat (left), with corresponding clear images (right). Texforms are generated using texture-synthesis algorithm from Deza et al. (2019) and preserve texture and form of original image while rendering image less recognizable. (B) **Example Trial**. On each trial, participants viewed a texform for 4 seconds while simultaneously settling on a best guess for what the original image was. Participants then rate how confident they are in their best guess on a continuous scale from 0 to 100, rate how curious they are to see the original image from 0 to 100, and finally see the original image for 2 seconds. (C) **Confidence predicts curiosity**. All variables were z-scored within participant, the black curve is the curve of best fit (fixed effects) from the polynomial regression model (*Methods*, Eq. 1). For visualization purposes only, points were binned according to confidence (30 bins: equal number of points in each bin). The gray error bars indicate mean curiosity +/- 1 SEM.

### Confidence and curiosity ratings share a negative, quadratic relationship

We found that perceptual curiosity and confidence ratings were negatively related (*Methods, Behavioral Analysis*): curiosity tended to peak at relatively low confidence and declined as individuals became more confident in their answer (**Fig. 1C**). The relationship had a significant quadratic component (mixed-effects model with linear and quadratic terms from Eq. 1; *β*_linear_ = -13.46, p < 0.0001, 95% CI = [-15.8 -11.0]; *β*_quadratic_ = -5.60, p < 0.0001, 95% CI = [-7.21 -3.99]) and model comparisons showed that models that contained both linear and quadratic terms were superior to those that contained only a linear term (BIC_quadratic_ - BIC_linear_=-180). Furthermore, curiosity was still related to confidence (*β*_linear_ = -13.46, p < 0.0001, 95% CI = [-15.8 -11.0]; *β*_quadratic_ = -5.58, p < 0.0001, 95% CI = [-7.19 -3.98]) even when including low-level visual properties (luminance, contrast, and spatial frequency; Eq 3 in *Methods*) as covariates and low-level visual properties were not reliable predictors of either curiosity (luminance (*β* = 0.70, p = 0.11, 95% CI = [-0.17 1.57]); contrast (*β* = -0.36, p = 0.41, 95% CI = [-1.2 0.51]); spatial frequency (*β* = -0.73, p = 0.16, 95% CI = [-1.75 0.30];) or confidence (luminance (*β* = -0.30, p = 0.60, 95% CI = [-1.44 0.84]); contrast (*β* = 1.20, p = 0.056, 95% CI = [-0.07 2.3]); spatial frequency (*β* = 0.76, p = 0.19, 95% CI = [-0.38 1.90];). This suggests that the relationship between confidence and curiosity is not contaminated by variation in low-level visual properties. Thus, consistent with epistemic curiosity (Kang et al. 2009; Gruber et al. 2014), perceptual curiosity and confidence showed a negative quadratic relationship, with curiosity being higher when participants had low or intermediate confidence in recognizing an item.

### OTC Certainty is positively correlated with confidence and negatively correlated with curiosity

To analyze the sensory underpinnings of perceptual curiosity, we focused on an anatomically-defined region of interest (ROI) in the occipitotemporal cortex (OTC), which encodes animal and man-made object categories across multi-voxel activity patterns (Kriegeskorte 2008; Konkle & Caramazza 2013) and is thus a good candidate area for representing multivariate certainty about the categories in our task.

Because existing measures of sensory certainty are limited to elementary visual features like orientation, and rely on strict assumptions about single-neuron tuning curves that are not compatible with the complex stimuli in our task, our first goal was to develop a measure of sensory certainty from multi-voxel activity patterns in OTC (“OTC Certainty”).

To this end, we ran a localizer task that was delivered unannounced after the main curiosity task (*Methods, Design & Procedure, Localizer Task*) in which participants viewed alternating miniblocks of undistorted animal and man-made object images (different from those in the main task). We then used a whole-brain general linear model to estimate the OTC multi-voxel activity pattern evoked by each miniblock (*Methods, fMRI Analysis, GLM #1*). We verified that activity patterns evoked by same-category miniblocks (i.e., animal/animal and man-made/man-made) showed higher pairwise correlations relative to patterns evoked by different-category miniblocks (i.e., animal/man-made; average Pearson’s r, 0.80 versus 0.58; p = 0.008), confirming that our OTC ROI conveys reliable category representations. We then averaged the responses to miniblocks of each category to obtain, respectively, an “animal template” and a “man-made template” -- the average multi-voxel activity pattern expected from images in each category.

Next, returning to the perceptual curiosity task, we measured the OTC activity pattern evoked by each texform (*Methods, fMRI Analysis, GLM #2*) and calculated the Pearson correlation coefficients between this activity pattern and each of our templates. Thus, for each texform-evoked activity pattern, r_a_ measures its correlation with the animal template and r_mm_ measures its correlation with the man-made template (**Fig 2B**). Texform-evoked activity patterns were more highly correlated with the matching relative to non-matching template (average Pearson r, 0.50 versus 0.43; paired t-test p = 0.01). Thus, the OTC responses generalized across the perceptual curiosity and localizer tasks and the coefficients r_a_ and r_mm_ were valid measures of the similarity between a texform-evoked pattern and the multivariate representation of the animal and man-made categories.

**Figure 2:**
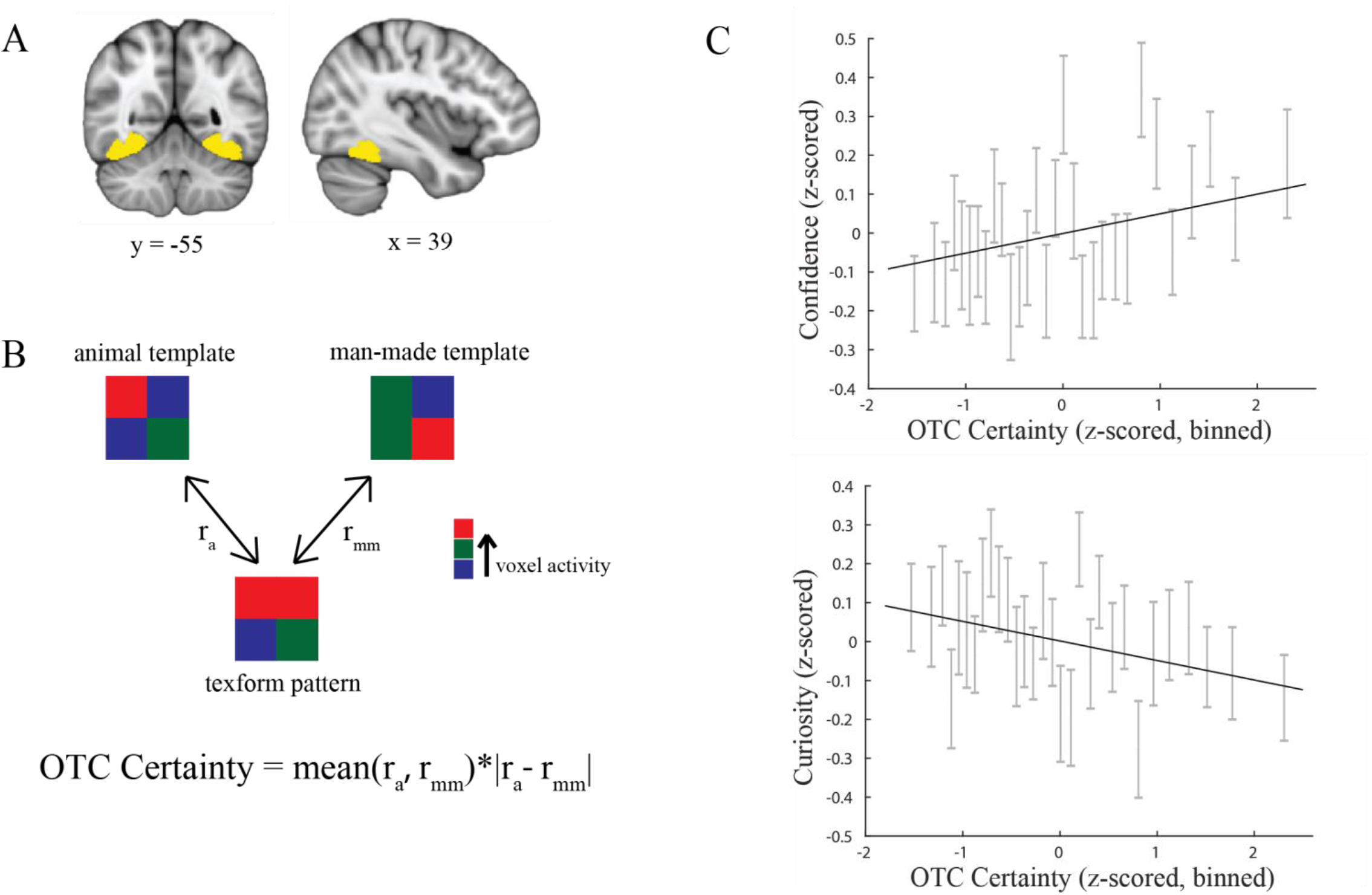
**(A) Anatomical OTC ROI**. Coronal (left), sagittal (right) views showing the OTC ROI used for subsequent analyses (yellow). **(B) Method for quantifying OTC Certainty**. We extracted category templates from an independent localizer by averaging multi-voxel patterns from animal and man-made miniblocks. We then correlated each category template with the multi-voxel pattern elicited by the texform on each trial and calculated OTC Certainty as the product of the average and absolute value of the difference in these two correlations (see text for details). **(C) OTC Certainty predicts confidence and curiosity**. All variables were z-scored within participant, the black line is the line of best fit (fixed effects) from the linear regression model (*Methods*, eq. 4 & 6). For visualization purposes only, points were binned according to OTC Certainty (30 bins: equal number of points in each bin). The gray error bars indicate mean confidence (top)/curiosity(bottom) and +/- 1 SEM in each OTC Certainty bin.

Next, motivated by the importance of capturing both model uncertainty and approximation uncertainty as emphasized in the machine learning literature (Huellermeir & Weigeman 2020); see *Discussion*), we defined a metric of OTC Certainty based on the average and relative values of r_a_ and r_mm_:

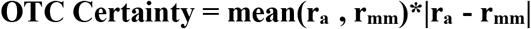

The first term in our function corresponds to model uncertainty – the degree to which a measured response is consistent with the hypotheses that are considered by the model. In our case, this is the extent to which a texform-evoked activity pattern is consistent with *either* of the templates, i.e., the mean of r_a_ and r_mm_). The second term in the function corresponds to approximation uncertainty – the degree to which a measured response distinguishes between the alternative hypotheses. In our case, this is the extent to which a texform-evoked activity pattern is more consistent with the animal *versus* the man-made template, i.e., the absolute difference between r_a_ and r_mm_.

Defined in this way, the metric of OTC Certainty is highest if a texform-evoked activity pattern is biased toward one category *and* is consistent with at least one of the categories we consider. For instance, for a fixed average coefficient of 0.3, OTC Certainty is higher if the difference between r_a_ and r_mm_ is higher (e.g., r_a_ = 0.1 and r_mm_= 0.5 vs r_a_ = 0.2 and r_mm_= 0.4). Thus, a low absolute difference between r_a_ and r_mm_ (the second term in the equation) reduces certainty because it suggests that the OTC does not clearly associate the texform-evoked pattern with one of the model categories. In addition, for a fixed difference of 0.2 between of r_a_ and r_mm_, the measure of OTC Certainty is higher if the average of the coefficients is higher (e.g., r_a_ = 0.3 and r_mm_= 0.5 vs r_a_ = 0.1 and r_mm_= 0.3). A low average coefficient (the first term in the equation) reduces certainty because it suggests that none of the categories we consider is a good model for the texform-evoked pattern.

We found that this metric of OTC Certainty had a positive relation with confidence ratings (*β* = 1.95, p = 0.0008, 95% CI = [0.80 3.09]; Eq. 4; **Fig. 2C top**) and a negative relation with curiosity ratings (*β* = -1.21, p = 0.007, 95% CI = [-2.08 -0.33]; Eq. 6; **Fig. 2C bottom**). Both relationships were linear, with model comparisons favoring models of confidence and curiosity that contained only linear terms for OTC Certainty over those that contained both linear and quadratic terms (confidence: BIC_quadratic_-BIC_linear_=17; curiosity: BIC_quadratic_-BIC_linear_=23). These relationships could not be explained by confounds related to head movements (which were included as nuisance regressors in *GLM #2; Methods*), cursor displacement (starting position was randomized on each trial), the scaling factor controlling image distortion (which was constant; *Methods*). Importantly, OTC univariate activity was not related to curiosity (*β* =-0.23, p=0.63, 95% CI = [-1.17 0.72]), suggesting that curiosity ratings are specifically associated with multivariate OTC Certainty rather than the univariate activity.

An important question is whether these associations between OTC certainty and curiosity reflect low-level image properties. For example, in their previous experience, people may have associated a low-level property like luminance or spatial frequency with a particular level of perceptual confidence and may have used this property, rather than OTC certainty, as the basis for their confidence and curiosity ratings (Geurts et al. 2022). To control for this possibility, we entered the luminance, contrast, and spatial frequency (*Methods*) of our texforms as covariates in the model linking OTC certainty and curiosity (*Methods*). OTC Certainty was related to curiosity even when including low-level visual properties as covariates (*β* = -1.15, p = 0.011, 95% CI = [-2.03 -0.26]) and low-level visual properties were not reliable predictors of OTC Certainty (luminance (*β* = 0.68, p = 0.12, 95% CI = [-1.07 0.68]); contrast (*β* = -0.19, p = 0.66, 95% CI = [-1.07 0.68]); spatial frequency (*β* = -0.71, p = 0.17, 95% CI = [-1.72 0.31]). Thus, the relationship between OTC Certainty and curiosity is not contaminated by variation in low-level visual properties.

In an additional analysis, we reasoned that, if our findings were explained by image-level confounds, activity in the primary visual cortex (V1) may show a behavioral relationship similar to that shown by the OTC. We thus replicated our analyses in V1, by first measuring multi-voxel activity pattern templates for animal and man-made objects in this area and then calculating the Pearson correlation coefficients between these templates and the texform-evoked activity patterns. In contrast to the OTC, texform-evoked activity patterns in V1 were not more correlated with the matching relative to non-matching template (average Pearson r, 0.57 versus 0.57; paired t-test p = 0.90), meaning that V1 did not differentiate animal and man-made object categories. This is a precursor for multivariate certainty, making it unlikely that multivoxel patterns in this area provide a meaningful signal for perceptual curiosity (and indeed, certainty derived from these patterns did not predict confidence (*β* = 0.12, p = 0.8, 95% CI = [-1.04. 1.30]) or curiosity (*β* = 0.04, p = 0.9, 95% CI = [-0.90, 0.99])). Thus, perceptual curiosity could not be explained by visual representations in V1, which suggests that higher order visual representations are important for evoking perceptual curiosity rather than lower-level visual representations.

### vmPFC, but not ACC, mediates the link between OTC and curiosity

Our finding that OTC Certainty was related to both curiosity and confidence (**Fig. 2**), together with the fact that the two ratings were correlated (**Fig. 1C**), raise questions about the link between OTC Certainty and curiosity. One scenario, suggested by previous studies of sensory certainty (van Bergen et al. 2020; Geurts et al. 2022), is that multivariate OTC Certainty is transformed into a univariate confidence representation elsewhere in the brain, which is in turn linked to curiosity. Alternatively, OTC Certainty may influence curiosity ratings independently of univariate confidence representations. To adjudicate between these possibilities, we analyzed whether two frontal brain areas implicated in perceptual confidence, the vmPFC (**Fig. 3A;** Mackey & Petrides Atlas) and ACC, (**Fig. 3B**; Harvard-Oxford Atlas), mediate the relationship between OTC Certainty and curiosity.

**Figure 3:**
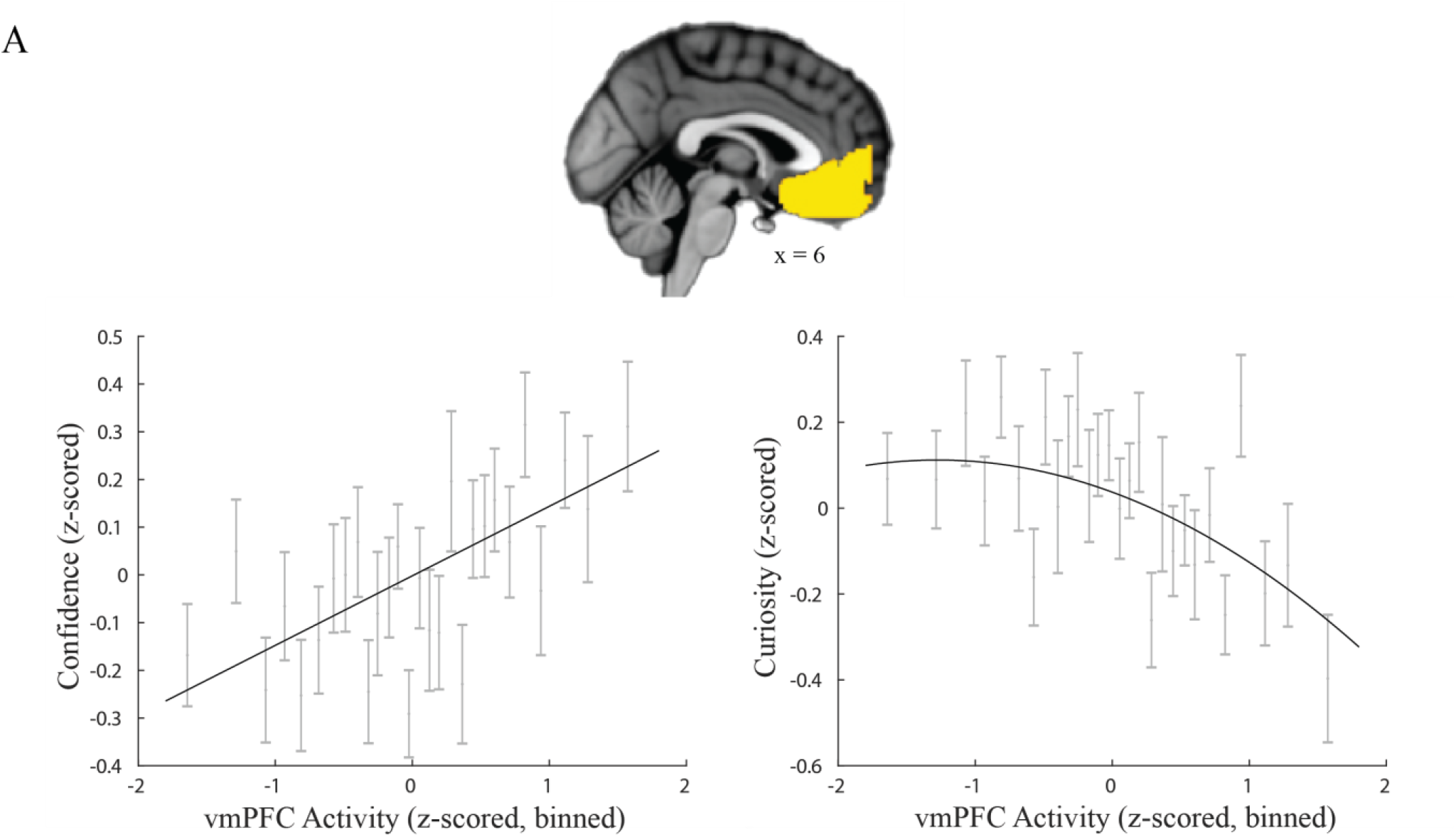
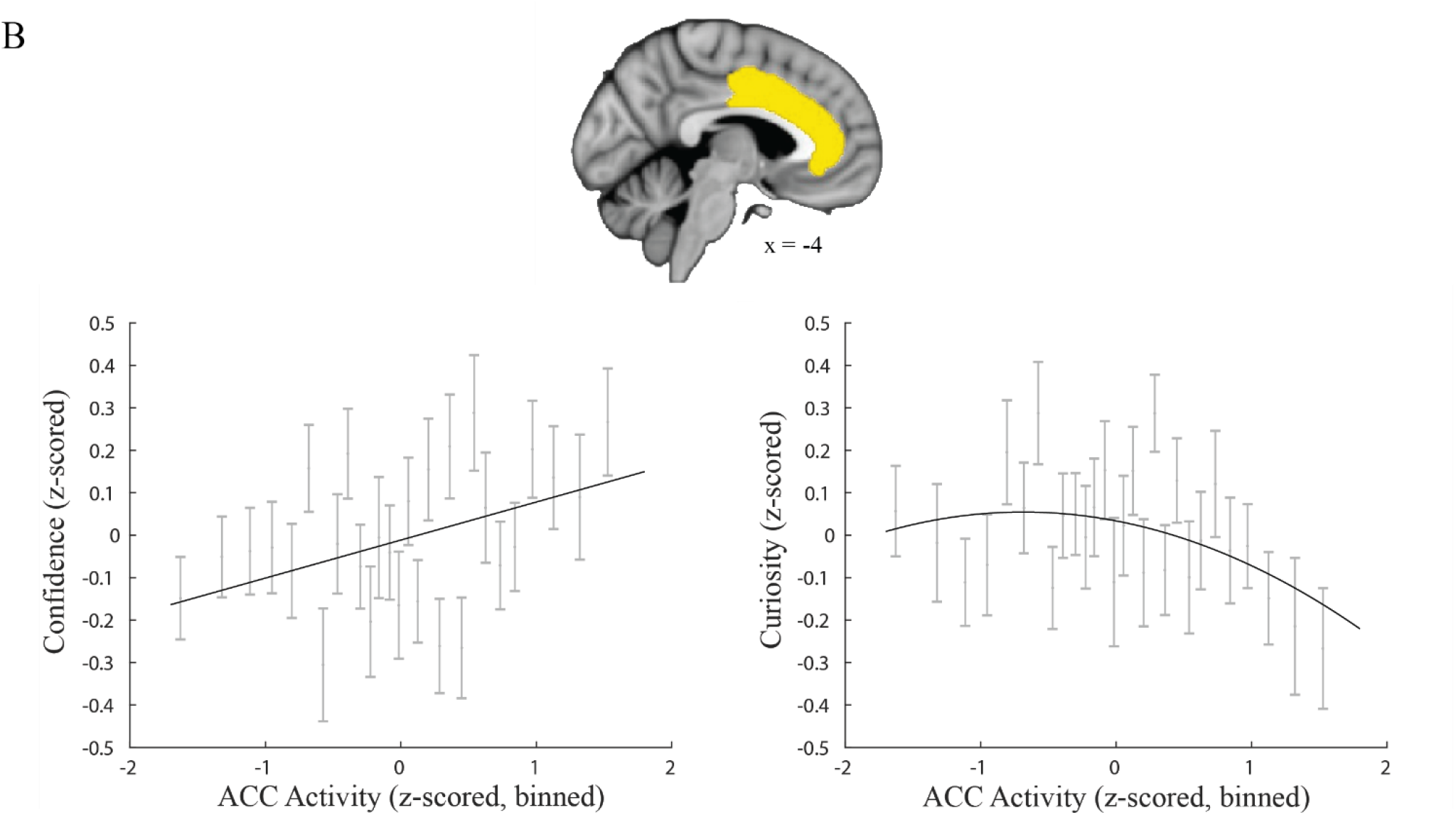
**(A) The vmPFC encodes curiosity and confidence**. The top panel shows the anatomical ROI for vmPFC (yellow). The bottom panels show the relationship between vmPFC activity and confidence and vmPFC activity and curiosity in the same format as in **Fig. 2C. (B) The ACC encodes curiosity and confidence**. The top panel shows the anatomical ROI for the ACC (yellow). The bottom panels show the relationship between ACC activity and confidence and ACC activity and curiosity in the same format as **Fig. 3A**.

Both areas showed univariate activity that scaled with confidence and curiosity (*Methods*). Consistent with previous studies (Gherman et al. 2018; Lebreton et al. 2015; Bang et al. 2018), confidence ratings were positively associated with univariate activity in the vmPFC (*β* = 3.32, p<0.0001, 95% CI = [1.80 4.85]; **Fig. 3A, left)** and the ACC (*β* = 2.34, p<0.0001, 95% CI = [0.88 3.79]; **Fig. 3B, left**). In contrast, curiosity ratings were negatively associated with univariate activity in both areas (vmPFC: *β*_linear_ = -2.43, p<0.0001, 95% CI = [-3.58 -1.28]; **Fig. 3A, right**; ACC: *β*_linear_ = -1.14, p<0.0001, 95% CI = [-2.26 -0.02]; **Fig. 3B, right**). In both areas, the relation between confidence and activity was linear (i.e., was better fit by models with only a linear term relative to models that also included a quadratic term; vmPFC: BIC_quadratic_ - BIC_linear_ =23; ACC: BIC_quadratic_ - BIC_linear_ =26). In contrast, curiosity ratings produced moderate evidence for an additional quadratic relationship. In the ACC, the relation between univariate activity and curiosity was equally well fit by models with and without a quadratic term (BIC_quadratic_- BIC_linear_=0; *β*_quadratic_ = -0.53, p=0.005, 95% CI = [-1.03 -0.02]) and in the vmPFC, the model that included a quadratic term produced a marginally better fit (BIC_quadratic_- BIC_linear_=-3; *β*_quadratic_ = -0.66, p = 0.005, 95% CI = [-1.13 -0.20]). Furthermore, activity in ACC and vmPFC was related to both confidence (ACC: *β* = 2.42, p = 0.0011, 95% CI = [0.96 3.88]; vmPFC: *β* = 3.37, p <0.0001, 95% CI = [1.84 4.90]) and curiosity (ACC: *β*_linear_ = -1.41, p = 0.017, 95% CI = [-2.58 -0.25]; *β*_quadratic_ = -0.54, p = 0.03, 95% CI = [-1.04 -0.03]; vmPFC: *β*_linear_ = -2.48, p <0.0001, 95% CI = [-3.63 -1.33]; *β*_quadratic_ = -0.66, p = 0.005, 95% CI = [-1.12 -0.20]) even when including low-level visual properties as covariates and low-level visual properties were not reliable predictors of either ACC (luminance (*β* = 1.34, p = 0.24, 95% CI = [-2.56 1.37]), contrast (*β* = -2.07, p = 0.06, 95% CI = [-4.29 0.15]), or spatial frequency (*β* = -2.00, p = 0.11, 95% CI = [-4.45 0.44]) or vmPFC activity (luminance (*β* = -0.61, p = 0.52, 95% CI = [-2.49 1.27], contrast (*β* = -0.90, p = 0.35, 95% CI = [-2.77 0.97], or spatial frequency (*β* = -2.05, p = 0.08, 95% CI = [-4.40 0.28])). Thus, the relationship between confidence/ curiosity and vmPFC/ACC activity is not contaminated by variation in low-level visual properties.

To understand the relationship between OTC Certainty, curiosity, and vmPFC and ACC, we performed mediation analyses (*Methods, fMRI Methods, Mediation Analysis)* comparing two models: a mediated model, in which vmPFC (or ACC) mediates the link between OTC Certainty and curiosity (**Fig 4A, top**), and an unmediated model, in which vmPFC (or ACC) and OTC Certainty independently contribute to curiosity (**Fig 4A, bottom**). We verified that vmPFC and ACC activity were positively and linearly associated with OTC Certainty, fulfilling a necessary condition for the mediation analysis (vmPFC: *β*_linear_ = 5.55, p < 0.0001, 95% CI = [2.91, 8.10])); BIC_quadratic_ - BIC_linear_ = 17; **Fig 4B, top**; ACC: *β*_linear_ = 9.00, p < 0.0001, 95% CI = [6.70, 11.3])); BIC_quadratic_ - BIC_linear_ = 3; **Fig 4B, bottom**).

**Figure 4:**
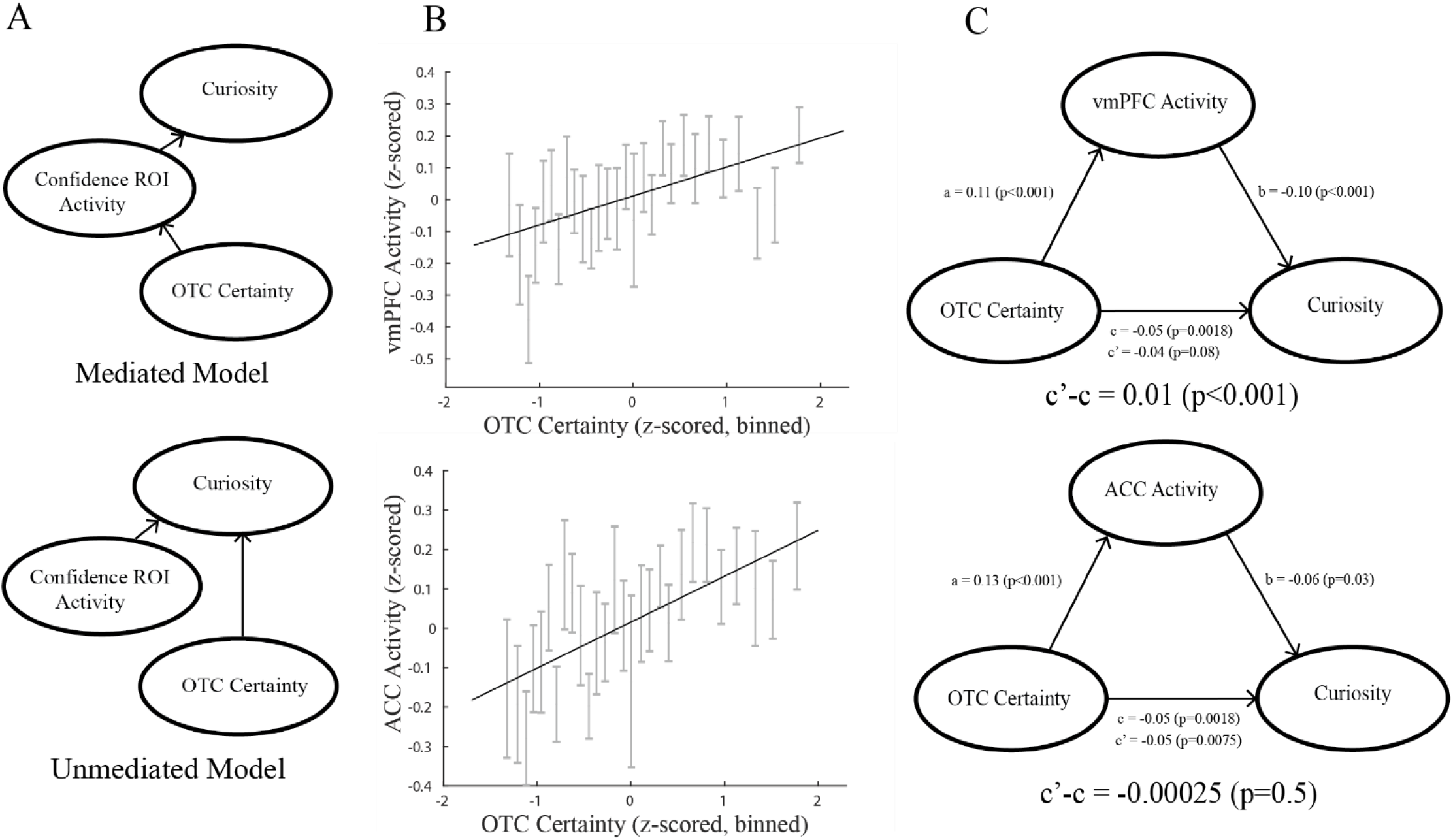
**(A) Candidate statistical models for the relationship between OTC certainty and curiosity**. In the “Mediated Model”, the relation between OTC certainty and curiosity ratings is mediated by activity in another area encoding univariate confidence (“confidence ROI”). In the “Unmediated Model”, OTC certainty and the confidence ROI independently predict curiosity. **(B) OTC certainty significantly correlates with vmPFC and ACC activity**, fulfilling a necessary condition for mediation analysis. Conventions as in Fig. 2C. **(C) Significant mediation occurs for the vmPFC (top) but not the ACC (bottom)**. In each path diagram, a, b, and c are the Pearson correlation coefficients between, respectively, OTC Certainty & the confidence ROI, the confidence ROI & curiosity, and OTC Certainty & curiosity respectively. c’ is the correlation coefficient between OTC Certainty and curiosity, when the confidence ROI response is controlled for.

According to the Baron & Kenny Mediation algorithm, mediation occurs if the effect of OTC Certainty on curiosity is reduced after controlling for activity in the confidence ROIs (vmPFC and ACC) (Baron & Kenny 1998). The results showed that vmPFC, but not ACC, significantly mediated the relationship between OTC Certainty and curiosity. In the model testing for vmPFC mediation, the link between OTC Certainty and curiosity showed a negative association that was significant before accounting for vmPFC activity (bootstrapped c, p < 0.001) but became weaker (closer to zero) after accounting for vmPFC activity (bootstrapped c’-c = 0.0109, two-tailed p < 0.001; **Fig 4C, top**). Thus, the vmPFC significantly mediated the relationship between OTC Certainty and curiosity, consistent with the mediated model (**Fig. 4A, top**). In contrast, in the model testing for mediation by the ACC, the link between OTC Certainty and curiosity showed a negative association that was significant before accounting for ACC activity (**Fig. 4C, bottom, c**, p < 0.001), and was unchanged after accounting for ACC activity (bootstrapped c’-c = 0, two-tailed p = 0.50; **Fig 4C, bottom**). Thus, the ACC did not mediate the relationship between OTC Certainty and curiosity, consistent with the unmediated model (**Fig. 4A, bottom**). Direct comparison confirmed that c’ (i.e. the effect of OTC Certainty on curiosity after accounting for activity in the confidence ROI) was significantly weaker in the vmPFC vs ACC model (Kolmogorov-Smirnov Test on bootstrapped c’ distributions, ks-stat = 0.60, p < 0.001). In conclusion, the vmPFC seems to play a specialized role in mediating the link between OTC Certainty and curiosity that is not shared by the ACC.

## Discussion

We tested the novel hypothesis that a neural representation of multivariate sensory certainty is transformed into a univariate confidence signal that generates curiosity. We used a new task in which participants rated their confidence and curiosity about images that were difficult to recognize and showed that, like curiosity about trivia questions (i.e. epistemic curiosity), perceptual curiosity had a negative, quadratic relation with confidence. By using synthetic images of animals and man-made objects (“texforms”) whose distortion was precisely controlled (Long et al. 2018), we could elicit multi-voxel categorical representations in the OTC and derive a trial-by-trial measure of the sensory uncertainty of this representation. We showed that OTC Certainty was negatively correlated with curiosity ratings. Moreover, this relationship was specifically mediated by the vmPFC but not the ACC, despite the fact that both areas provided univariate responses to confidence and the ACC had been associated with perceptual curiosity (Jepma et al. 2012). These results go beyond previous studies that identified brain regions encoding curiosity about trivia questions (Kang et al. 2009; Gruber et al. 2014) or presented blurry visual images without examining sensory representations or collecting curiosity and confidence ratings (Jepma et al. 2012), and suggest a mechanism by which a neural representation of the (un)certainty about an event can generate curiosity.

The hypothesis that sensory areas provide a representation of stimulus certainty that is read out into a representation of confidence has been tested in human V1 (Geurts et al. 2022). The study derived a generative model for decoding multivariate certainty that was tailored to elementary features and relied on the assumption that V1 cells have cosine-tuning for orientation (vanBergen et al. 2021; Geurts et al. 2022). Although this assumption is well-supported for early visual areas (e.g., Walker et al. 2020) it does not apply to higher-level visual regions like the OTC, where individual neurons and voxels show complex selectivity to stimulus categories rather than elementary features (Kriegeskorte et al. 2008; Thorat et al. 2019).

When devising our metric of OTC Certainty, therefore, we could not derive algorithms for decoding certainty starting from first principles of cosine tuning as was done by Geurts and colleagues. Instead, we relied on the view that the certainty of an analytical model stems from two synergistic components. Model certainty captures the extent to which the data fall within the space of hypotheses that are considered by the model; in addition, approximation certainty captures how well the model can differentiate between the hypotheses (Hullermeier & Waegeman 2022). We verified that, in our data, the correlation coefficients r_a_ and r_m_ estimated the evidence in favor of the modeled categories (i.e., scaled with the true category of a texform). Therefore, the average of r_a_ and r_m_ captured model certainty (whether the activity pattern was consistent with at least one of the model categories) and the absolute difference between r_a_ and r_m_ measured approximation uncertainty (the distinctiveness of the representations within the hypothesis space). Our combined OTC Certainty metric was thus highest if a texform-evoked activity pattern was biased toward one category and was consistent with at least one of the categories. The fact that OTC Certainty correlated with subjective ratings of curiosity and confidence suggests that the idea that visual areas encode distributed representations of certainty can be extended to higher-order visual areas. Moreover, the results provide a new method for measuring certainty that avoids strong assumptions about sensory tuning and thus may generalize to other domains beyond visual features – including, potentially, the encoding of semantic concepts or categories.

A potential limitation of our approach is that multivariate certainty was measured at the level of animal and man-made object categories, while participants were instructed to generate best guesses about the distorted image with as much specificity as they could and may have generated guesses at the level of exemplars. However, our approach requires a single assumption: that generating a guess at any level of specificity within one category activates patterns elicited by that category more than patterns elicited by other categories. As we show, this assumption was met in our study, establishing the validity of our approach. Nevertheless, future studies can attempt to quantify multivariate certainty at the level of exemplars rather than categories and determine whether and how more granular representations account for the elicitation of curiosity.

Curiosity is considered to be a motivating drive to acquire information even if it comes with no monetary rewards. Although curiosity is non-instrumental, and we linked vmPFC/ACC activity to curiosity, other studies indicate that vmPFC (Kable & Glimcher 2007: Clithero & Rangel 2014) and ACC (Goh et al. 2021; Cai & Padoa-Schioppa 2012; Yee et al. 2021) have roles in representing subjective value. This raises the possibility that activity in vmPFC/ACC in our study may be representing participants’ subjective value of seeing the undistorted image. Indeed, according to prominent theories of curiosity, high curiosity states may be associated with high subjective value (Litman 2005; Gruber & Ranganath, 2020; Kang et al. 2009). However, our results pose a problem for this interpretation of vmPFC/ACC activity. In our study, activity in these areas was highest when curiosity was relatively low, and hence, when subjective value may have been relatively low as well, inconsistent with the positive relationship between vmPFC/ACC activity and subjective value that is reported in the literature, whereby activity tends to be highest when subjective value is high (Kable & Glimcher, 2007; Shenhav et al. 2013; Yee et al. 2021). Thus, the confidence and curiosity relationships we observed with vmPFC/ACC activity are likely separable from signals of subjective value.

Despite both vmPFC and ACC having positive relationships with OTC Certainty and negative relationships with curiosity, we show that only the vmPFC mediated the relationship between OTC Certainty and curiosity. This result is consistent with the fact that the vmPFC, but not ACC, has direct anatomical connections with the OTC (Catanati 2005; Furl 2015). Moreover, the vmPFC encodes visual confidence in multiple tasks that involve both the discrimination of simple visual features (Gherman et al. 2018) and more complex tasks like retrieval from memory (Hebscher et al. 2016) and judgments about the pleasantness and age of a painting (Lebreton et al. 2015). Thus, our findings suggest that the vmPFC is in an ideal position to mediate links between OTC Certainty and downstream behavioral judgements of curiosity.

In contrast, the fact that our results provide lower evidence for ACC mediation has several possible interpretations. One possibility is that the ACC computes confidence based on sources beyond OTC Certainty that have less impact on perceptual curiosity – e.g., factors related to response heuristics or biases (Maniscalco et al. 2016; Peters et al. 2017; Zylverberg et al. 2012; Merkel et al.) or elementary visual features encoded in V1 (Geurts et al. 2022). An alternative, not mutually exclusive, possibility is that the ACC is not primarily involved in generating curiosity based on certainty, but rather in recruiting cognitive functions to satisfy curiosity. This hypothesis is consistent with the role of ACC in executive function (Shenhav et al 2013) and its hypothesized role in conveying signals of confidence and certainty to the fronto-parietal network to implement attentional orienting for information gathering (Silvetti et al., 2023; Gottlieb et al. 2021; Foley et al 2017; Horan et al 2020; White et al. 2019). A final possibility is that the ACC plays distinct roles in instrumental decisions, when information is used to guide actions that lead to rewards, versus non-instrumental decisions like curiosity, which are intrinsically motivated and independent of external rewards (Gruber et al. 2014; Gottlieb et al. 2021). Thus, the specific role of the ACC in various types of decisions involving confidence and the demand for information remains an important topic for future investigations.

Our finding that the vmPFC mediates the relationship between OTC Certainty and curiosity is consistent with previous work on the relation between certainty and confidence (Meyniel et al. 2015; Pouget et al. 2016; Geurts et al 2022) and, more broadly, with the representational untangling hypothesis (DiCarlo & Cox 2007; Russo et al. 2018) whereby multivariate neural representations are transformed into lower dimensional representations to generate behavior. However, one unexpected result in our study is that curiosity ratings showed a linear rather than quadratic relationship with OTC Certainty. Given our findings that the vmPFC mediates the link between OTC Certainty and curiosity and shows a quadratic relationship to curiosity, one might expect a similar quadratic relationship between OTC Certainty and curiosity. One possibility is that we failed to detect a quadratic relationship between OTC Certainty and curiosity due to statistical noise (especially since the evidence for a quadratic relationship for vmPFC was moderate). A second, more intriguing possibility is that there is an additional pathway that mediates the link between OTC Certainty and curiosity in which all the relationships are linear. Future work can examine whether these pathways exist and how they uniquely contribute to curiosity.

In conclusion, our results are consistent with the hypothesis that multivariate representations of certainty are transformed into univariate confidence in frontal cortex to generate curiosity. We speculate that this transformation might generalize outside of visual processing into other stimulus domains in which stimulus representations are probabilistic (Pouget et al. 2015; Lindskog et al. 2021) and thus are likely described by multivariate certainty (Menyiel et al. 2015). Together, our findings demonstrate the neural mechanisms of curiosity by explaining how a neural representation of an event can give rise to curiosity about that event.

## Methods

### Participants

Thirty-two individuals (17 female; 18–35 years old; all right-handed; all normal or corrected-to-normal vision) participated for monetary compensation ($20/hour; $40 in total). The study was approved by the Institutional Review Board at Columbia University. Participants were recruited via mailing lists and a research participation pool at Columbia University. Written informed consent was obtained from all participants. Participants also passed a health and safety screening on their eligibility for the MRI scanner.

### Stimuli

To elicit curiosity, we used texforms (Konkle et al 2018; Deza et al. 2019). We first collected 42 images of animals and 42 images of man-made objects from the Konkle Lab image database and normalized them for contrast and luminance across the whole set using the SHINE Toolbox in MATLAB. Then, we used an existing algorithm (Deza et al. 2019) that calculated thousands of first- and second-order image statistics from individual pooling regions overlaid across each image. Finally, we generated the texform by starting from a white noise display and coercing it to match the measured image statistics using stochastic gradient descent (100 iterations). The resulting texform looked like a distorted version of the original one. The size of the pooling regions (spatial pooling factor) determined the degree of distortion; all the images we used had a constant pooling factor of 0.28.

To investigate if low-level visual properties can predict our variables of interest (e.g. OTC Certainty), we measured the luminance, contrast, and spatial frequency of the texforms. Luminance was determined by calculating the mean of the intensity histogram, while root-mean-squared (RMS) contrast (i.e. standard deviation of the luminance distribution) was used to measure contrast, as is typical in vision studies using natural stimuli (Peli 1990). Spatial frequency was calculated by first using 2-D Fast Fourier Transform (fft2 in Matlab), calculating power spectra in each dimension, and calculating the square root of the sum of squares of the resulting power spectra (i.e. sqrt(power(x)^2+power(y)^2)). Average spatial frequency is then calculated as the slope of this resulting power spectrum, and provides a measure of the amount of high vs low frequency information in the image. This approach to calculate spatial frequency content has been used in previous studies (Li et al. 2001; Eskicioglu & Fischer 1995; Flitcroft et al. 2020).

### Design & Procedure

#### Perceptual Curiosity Task

Stimuli were presented using the Psychophysics Toolbox for MATLAB (Psychtoolbox). For all trial components that required participants to enter a rating, participants did so on a continuous scale from 0-100 using an MR-compatible trackball. The initial slider position was randomized on every trial. Participants had up to 5 seconds to respond to each prompt; the trial advanced to the next screen after a response was entered.

The design of the perceptual curiosity task was inspired by Gruber et al. (2014) and Jepma et al. (2012). On each of 84 trials, participants viewed a texform (see *Stimuli*) of either an animal or a man-made object, which remained on the screen for 4s. While viewing the texform, participants were instructed to come up with their *best guess* for what the original (undistorted) image was. Next, participants are prompted to rate their confidence in their best guess for the original image and their curiosity to see the original image. Finally, participants viewed the original image for 2s. Trials were divided evenly into four runs. Importantly, participants were paid a fixed amount of $40 regardless of performance. Thus, their confidence and curiosity ratings are independent of monetary incentives.

#### Localizer Task

After this task, participants completed an unannounced localizer run, in which they viewed alternating miniblocks of clear (undistorted) animal and man-made object images that were not seen earlier in the task. Each of the 24 miniblocks (12 animal miniblocks and 12 man-made miniblocks) consisted of the presentation of 20 images presented in rapid succession (333 ms per image and 333 ms inter-stimulus interval). Between miniblocks, participants were presented with a fixation screen (13 seconds), which allowed for separation of BOLD activity between miniblocks. During image presentation, participants completed a 1-back cover task, in which they were asked to detect and respond to repeat images using a button box.

### Behavioral Analysis

All mixed-effects models of behavior were conducted using the *fitlme* function in MATLAB.

#### Mixed-effect modeling of curiosity

To examine the relation between confidence and curiosity ratings, we constructed two mixed-effects models. In the Quadratic Model, curiosity was predicted by confidence, confidence^2^, and participant-specific random slopes (confidence|participant and confidence^2^|participant) and intercepts (1|participant).

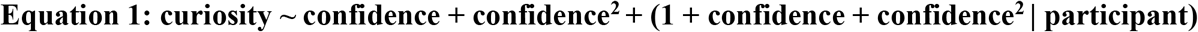

In the Linear Model, curiosity was predicted by confidence and participant-specific random slopes (confidence|participant) and intercepts (1|participant).

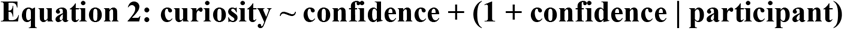

To examine whether the relation between confidence and curiosity persists when including low-level visual properties as covariates, we constructed a mixed-effect model in which curiosity was predicted from confidence, luminance, contrast, and spatial frequency, as well as participant-specific random slopes (confidence|participant, luminance|participant, contrast|participant, and spatial frequency|participant) and intercepts (1|participant).

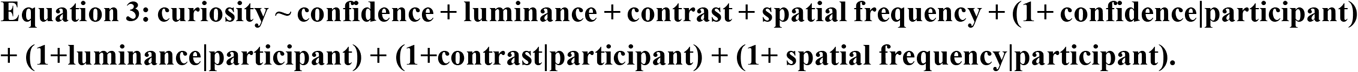

Independent variables across both models were z-scored within participant.

### MRI Acquisition

MRI data were collected on a 3T Siemens Magnetom Prisma scanner with a 64-channel head coil. Functional images were obtained with a multiband echo-planar image (EPI) sequence (repetition time = 2s, echo time = 30 ms, flip angle = 80°, acceleration factor = 3, voxel size = 2 mm isotropic; phase encoding direction: posterior to anterior), with 69 axial slices (14° transverse to coronal) acquired in an interleaved fashion. There were five functional runs: four for the perceptual curiosity task and one for the localizer task. Whole brain high resolution (1.0 mm isotropic) T1-weighted structural images were acquired with a magnetization-prepared rapid acquisition gradient-echo (MPRAGE) sequence.

### fMRI Analysis

#### Software

Preprocessing and analyses were performed using FEAT, FNIRT, and command-line functions in FSL (e.g., fslmaths). Subsequent analyses were performed using custom MATLAB scripts. Code available upon request.

#### ROI Definition

The vmPFC ROI was based on Mackey and Petrides (2014), but voxels were removed that overlapped with the corpus callosum. The OTC ROI and ACC ROI were derived from the Harvard-Oxford Brain Atlas using the atlas tool (threshold = 50) in fslview. All ROIs were in 1.0 mm MNI space.

#### Preprocessing

We performed brain extraction (using BET), motion correction (using MCFLIRT in FEAT), high-pass filtering (cut-off = 100 ms), and spatial smoothing (3mm FWHM Gaussian kernel). Functional images were registered to the standard 1mm MNI152 structural image using a non-linear transformation with 12 degrees of freedom. This registration was done on first-level analysis output (beta maps from individual trials or miniblocks).

#### GLM #1– Localizer GLM

We implemented a GLM for each participant that predicted BOLD activity of every voxel from a design matrix that modeled each localizer miniblock as a boxcar from the onset of the first image in each miniblock to the offset of the last image in each miniblock (16.66 s). Thus, for each of the 24 miniblocks, we included one regressor in the design matrix, yielding one beta map for each miniblock. This design matrix also contained fixed-body motion-realignment regressors (x, y, z, pitch, roll, and yaw) and their respective first derivatives. All regressors were convolved with a double-gamma hemodynamic response function. Autocorrelations in the time series were corrected with FILM pre-whitening.

#### Formation of animal and man-made object templates from GLM#1

To create the animal template, beta maps for the 12 animal miniblock regressors (from GLM #1) were averaged within the OTC ROI. To create the man-made object template, beta maps for the 12 man-made object miniblock regressors (from GLM #1) were averaged within the OTC ROI.

#### GLM #2– Single trial GLM

We implemented a GLM for each participant that predicted BOLD activity of every voxel from a design matrix that separately modeled the texform presentation on each trial as a 4s boxcar (i.e., 21 texform regressors per run). This design matrix also contained fixed-body motion-realignment regressors (x, y, z, pitch, roll, and yaw) and their respective first derivatives. Other regressors modeled trial components that were not of primary interest for fMRI analyses: one regressor that modeled all the confidence ratings periods, each as a boxcar with a duration equal to the participant’s response time (RT); one regressor that modeled all the curiosity ratings, each as a boxcar with a duration equal to the participant’s RT; and one regressor that modeled all the clear image presentations, each as a boxcar with a duration of 2 s. Each run was modeled separately, resulting in four different models per participant. All regressors were convolved with a double-gamma hemodynamic response function. Autocorrelations in the time series were corrected with FILM pre-whitening.

Univariate activity in vmPFC, ACC, and OTC during the texform period was obtained by averaging beta values across voxels, separately for each texform presentation. Multivariate activity patterns in OTC during the texform period were obtained by extracting the activity pattern across voxels, separately for each texform presentation.

#### Quantification of OTC Certainty

We calculated r_a_, the Pearson correlation coefficient between the animal template and texform pattern on a given trial, and r_mm_, the Pearson correlation coefficient between the man-made template and texform pattern. We quantified OTC Certainty on every trial as the product of two terms:

1. *relative evidence*, the absolute value of the difference between r_a_ and r_mm_
2. *mean evidence*, the mean of r_a_ and r_mm_

#### Mixed-effect modeling

We constructed mixed-effects models using the *fitlme* function in MATLAB to examine relationships between brain measures and between brain measures and behavior:

To examine the relation between confidence and OTC Certainty, we constructed two mixed-effects models. In the Quadratic Model, confidence was predicted by OTC Certainty, OTC Certainty^2^, and participant-specific random slopes and intercepts.

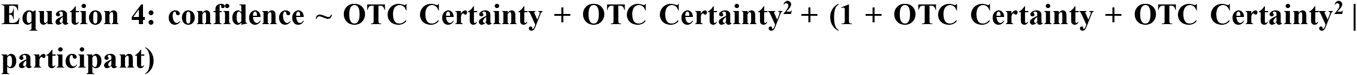

In the Linear Model, confidence was predicted by OTC Certainty and participant-specific random slopes and intercepts.

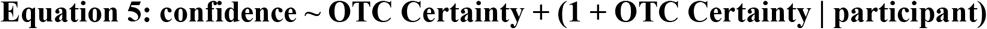

To examine the relation between curiosity and OTC Certainty, we constructed two mixed-effects models. In the Quadratic Model, curiosity was predicted by OTC Certainty, OTC Certainty^2^, and participant-specific random slopes and intercepts.

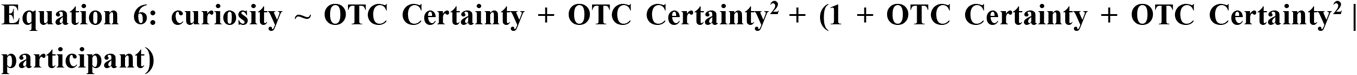

In the Linear Model, curiosity was predicted by OTC Certainty and participant-specific random slopes and intercepts.

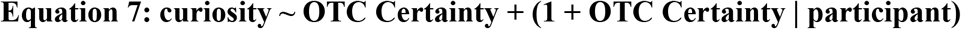

Similar models were constructed to examine the relationships between confidence and vmPFC activity, curiosity and vmPFC activity, OTC Certainty and vmPFC activity, confidence and ACC activity, curiosity and ACC activity, and OTC Certainty and ACC activity.

To examine whether the relation between OTC Certainty and curiosity persists when including low-level visual properties as covariates, we constructed a mixed-effect model in which curiosity was predicted from OTC Certainty, luminance, contrast, and spatial frequency, as well as participant-specific random slopes (curiosity|participant, luminance|participant, contrast|participant, and spatial frequency|participant) and intercepts (1|participant). Similar models were constructed to examine whether vmPFC/ACC activity could be predicted by low-level visual properties.

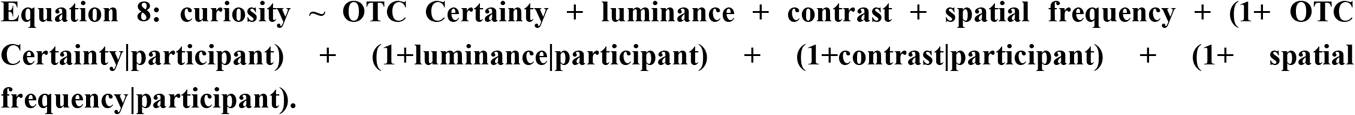

Independent variables across all models were z-scored within participant.

#### Model Comparison

We compared the relative goodness-of-fit of the Quadratic vs. Linear mixed-effect models using the Bayesian Information Criterion (BIC). Using the conventions from Rafferty et al. 2010, a BIC difference of >=2 indicates evidence for one model over another.

#### Mediation Analysis

To examine the hypothesis that univariate activity in our confidence ROIs mediates the correlation between OTC Certainty and curiosity, we used the Baron & Kenny approach to mediation (Baron and Kenny 1998) implemented in MATLAB by Wager (2009). This approach compares c’ (e.g., the effect size of the linear term of OTC Certainty on curiosity when controlling for confidence ROI activity) with c (e.g., the effect size of the linear term of OTC Certainty on curiosity alone). Statistical mediation occurs when c’ is smaller in magnitude than c. We included quadratic terms in all pairwise comparisons because some had a significant quadratic relationship; but we only examined the change of magnitude of the linear terms when testing for mediation, consistent with established mediation approaches. Similar results were obtained if we only incorporated linear terms in the mediation analysis. We z-scored the three variables of interest (OTC certainty, confidence ROI activity, and curiosity ratings) within participant, and bootstrapped with replacement to obtain a distribution of c’ and c values across 1000 iterations. We then took the difference of the c and c’ distributions and calculated the fraction of this new distribution that did not exceed 0. We then doubled this value to obtain our two-tailed p-value.

## Acknowledgments

The research described in this paper was supported by the National Institute of Mental Health as part of the National Research Service Award (Grant #:1F31MH125589), and the Zuckerman Institute MR Seed Grant Award (Grant #: CU-ZI-MR-S-0017) both awarded to Michael Cohanpour. We thank the Alyssano Group, Gottlieb Lab, Kriegeskorte Lab, Christopher Baldassano, Janet Metcalfe, and Yasmine El-Shamayleh for their valuable insight on this project; Ray Lee and Noreen Violante for their technical support with the MRI scanner; and Serra Favila, Heiko Schütt, and Javier Domínguez Zamora for their crucial revisions to the manuscript.

